# Modeling the human vaginal microbiome and its protection against pathogens using the replicator framework for invasion

**DOI:** 10.64898/2026.02.20.707042

**Authors:** Tomás Ferreira Amaro Freire, Marina Garcia-Romero, Erida Gjini

## Abstract

The human vaginal microbiota plays a central role in protecting against urogenital infections, including bacterial vaginosis, yeast infections, HIV, and urinary tract infections. However, the ecological mechanisms connecting microbial community structure to clinical indicators such as Nugent scores remain poorly understood. Although machine-learning approaches can accurately predict bacterial vaginosis (BV) from microbiota profiles, they provide limited biological insight. Here, we introduce a mechanistic framework that links vaginal microbiota composition to Nugent score and identifies key ecological interactions underlying health and disease-associated community states. We analyzed microbiota data from an already published North American women cohort, aggregating taxa at the phylum level to improve stability and interpretability. Using a replicator model, we quantified both the direct contributions of individual phyla and nonlinear effects arising from pairwise interactions, capturing transitions among healthy, intermediate, and BV-positive states. The model predicted BV-positive status with 92% accuracy for the 394 women in the study, *on par* with machine-learning benchmarks. More importantly, it provides a clear ecological interpretation of BV-associated community change. Beyond the BV setting, the model itself illustrates a general *proof-of-concept* for microbiota–invader links via the replicator formalism.

## Introduction

The human vaginal microbiota (VMB) plays a key role in protecting against urinary tract infections [1], pelvic inflammatory disease [2, 3], risk of miscarriage and pre-term labor [4], sexually transmitted infections [5, 6], and increased risk of HIV infection and transmission [7–9]. The VMB composition of reproductive-age cisgender women is dominated in most cases by *Lactobacillus* species, which produce lactic acid that helps maintain a low vaginal pH, creating an environment that is inhospitable to pathogens [10–12]. In contrast, shifts to communities more abundant in anaerobic bacteria such as *Gardnerella, Prevotella, Megasphaera, Atopobium, Sneathia*, and *Peptoniphilus*, compromise this protective environment and are associated with adverse clinical outcomes [2]. This polybacterial community shift is characteristic of bacterial vaginosis (BV), which is typically diagnosed in clinical settings using the Amsel criteria [13], while in research settings the Nugent score [14] is often used. The Nugent score ranges from 0 to 10 and is obtained by Gram-staining vaginal swabs, and microscopically assessing the relative abundance of three bacterial morphotypes: large Gram-positive rods (*Lactobacillus* morphotypes), small Gram-negative to Gram-variable rods (*Gardnerella vaginalis* and *Bacteroides* morphotypes), and curved Gram-negative or Gram-variable rods (*Mobiluncus* morphotypes). Morphotypes are scored semiquantitatively, such that decreased *Lactobacillus* and increased *Gardnerella*/*Bacteroides* and *Mobiluncus* morphotypes raise the total score. Samples are classified as healthy (0–3), intermediate (4–6), or BV positive (7–10). As the method depends on manual microscopic classification, Nugent scoring is subject to observer interpretation and inter-observer variability [15]. Although it is a useful clinical indicator for BV diagnosis in asymptomatic women, a better understanding of how it relates to ecological processes and microbial interactions governing transitions between different VMB health states, could lead to improvement of diagnosis and treatment.

The pioneering sequencing study by Ravel et al. [16, 17] established strong associations between VMB community composition and Nugent-defined health states, identifying five community state types (CSTs). Four CSTs were dominated by distinct Lactobacillus species, namely *L. crispatus, L. gasseri, L. iners*, and *L. jensenii*, while the fifth was characterized by high diversity and predominance of obligate anaerobic bacteria. More recent machine-learning studies have successfully predicted BV status from microbial profiles [18–21], but they offer limited insight into the ecological interactions and processes that drive these health-associated transitions.

Ecological dynamical models provide a complementary path toward mechanistic understanding. By representing the microbiota as systems shaped by interactions among taxa, such models have the potential to clarify key collective properties of community stability, invasibility, and resilience. The replicator equation [22, 23] stands out for offering a natural framework to model frequency-dependent dynamics and interactions within microbial consortia [24, 25]. These types of dynamical approaches, however, have rarely been applied to vaginal microbiome data, which has been mainly addressed via bioinformatics or machine learning algorithms. Neither have they been used to directly link clinically important indicators such as the Nugent score to microbiota compositions. Here, we bridge this gap, by introducing a simple and interpretable ecological model that relates vaginal microbiota structure to Nugent score categories. Proposing a replicator-dynamics framework, we analyze the dataset of the North American cohort of 394 women from Ravel et al. [16], to quantify and illustrate how phylum-level groups and their pairwise interactions contribute to defining healthy, intermediate, and BV-positive community states. This approach provides a new mechanistic framework for interpreting the ecological roles of major vaginal microbiota taxa, while generating quantitative and testable predictions about vaginal health and its conservation.

## Methods

### Data and data processing

The dataset comprises 394 vaginal microbiota profiles obtained from [16], along with their corresponding Nugent scores. Most samples fell in the 0–3 range (248/394), followed by the 7–10 range (97/394) and the 4–6 range (49/394). Across ethnicities, low scores (0–3) were most common in Asian (69/96) and White (77/97) participants, while higher scores (7–10) were more frequent among Black (42/104) and Hispanic (32/97) participants. Metagenomic classification was performed at the phylum level (see SI text S1 for details). In contrast to the original study, which assigned sequences to genus- and species-level taxa [16], we restricted the analysis to higher taxonomic ranks to reduce dimensionality while preserving ecologically meaningful community structure. Across all samples, 10 bacterial phyla were identified (see Fig. S2 for a summary of taxon frequencies and Table S1 for a list of common taxa belonging to each phylum): *Actinomycetota, Bacillota_A_368345, Bacillota_C, Bacillota_I, Bacteroidota, Campylobacterota_A, Fusobacteriota, Patescibacteria, Pseudomonadota*, and *Synergistota*. From these, *Synergistota* and *Campylobacterota_A* were present in low amounts (always below 5%). Vaginal *pH* and ethnic group were excluded from model training. Model performance was evaluated on the full cohort as well as within individual ethnic groups.

### Modeling Framework

To predict vaginal health status from vaginal microbiota composition, we use replicator dynamics, a mathematical framework originating from evolutionary game theory [22, 23]. The replicator equation lends itself naturally to the study of frequency-dependent invasion dynamics, a community’s invasion resistance [24], and has been recently applied to a gut microbiota dataset to explain invasion differences between strains of *E*.*coli* in mice [25]. This approach is particularly suitable for sequencing-based microbiome data (e.g., 16S rRNA gene sequencing), which provide relative abundances rather than absolute counts. Replicator dynamics naturally describe how species frequencies change over time and how species interactions shape this evolution. In this framework, the dynamics of a community of *N* species are modeled by an *N*-dimensional system of equations that describe the temporal evolution of the relative frequency of each species (*i* = 1,..*N*):

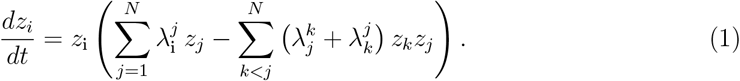

In this system, *Σ*_*i*_ *z*_*i*_ = 1 and 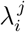 represents the pairwise invasion fitness of species *i* vs *j*, namely its initial growth rate into an equilibrium consisting only of species *j* [26], defined clearly for different species *i* ≠ *j*, forming together the invasion fitness matrix Λ. The diagonal of this matrix consists of zeros. Notice that the pairwise invasion fitness matrix does not uniquely determine underlying interactions between species. Many types of microscale interactions can lead to the same emergent Λ matrix [24, 27, 28], and it is only this net matrix that is ‘seen’ by the replicator, ultimately it is this matrix that governs all species dynamics.

The quadratic term in the replicator 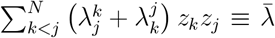 captures the average invasion resistance of the system [24]. The more positive this quantity is, the harder it is for the invader to grow, and vice versa. In this multispecies ecosystem, the initial growth of any newcomer at low frequency is given by:

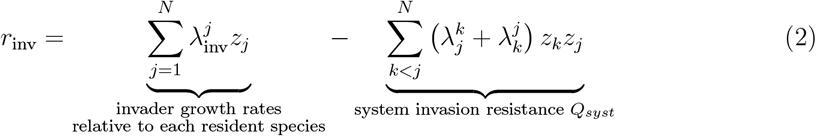

capturing the ability of a new species to invade the community when rare. Part of this initial growth rate, *r*_inv_, depends on its pairwise invasion fitness relative to resident species (linear term), as well as on the system’s mean invasion resistance (quadratic term, independent of the invader itself).

As a whole, *r*_inv_ indicates a local fitness, representing a first step for the invader to establish within a given resident multispecies environment. The higher and more positive this growth rate is, the easier it is for the outsider species to invade the resident community, indicating a disease-prone state. In contrast, the lower or more negative this rate is, the harder it is for the invader to succeed, indicating a healthy state.

On a very abstract level, unealthy (BV) and healthy states in vaginal microbiomes can be treated as two extremes on a spectrum of *r*_*inv*_ for a typical invader of the vaginal ecosystem. This will be our guiding analogy in the model framework, treating *r*_*inv*_ as our mathematical proxy for BV risk / vaginal health.

### From Nugent scores to invader growth rates

Because the original data do not measure invader growth rates, but isntead provide Nugent scores as summary indicators of women vaginal health (i.e. susceptibility to bacterial vaginosis), our first goal is to relate these empirical Nugent scores to the growth rate of an *invader* within the replicator framework. Translating a clinical, discrete diagnostic metric such as the Nugent score into a quantity with a clear ecological meaning requires an interpretation of what the *invader* represents.

Because high Nugent scores can arise from the proliferation of multiple anaerobic species, such as *Gardnerella, Prevotella, Megasphaera, Roseburia* and *Atopobium* [29], we interpret the *invader* more broadly as the collective ecological entities responsible for unhealthy states, pathogenic infection, vaginosis, and the like, i.e. a generic potential enemy capable of generating vaginal disease. In this sense, *r*_*inv*_ symbolically mirrors the system’s susceptibility to pathogenic invasion, or how easily the healthy state can be destabilized by invading taxa associated with BV. To connect this biological interpretation to the replicator framework, we map the discrete Nugent scale (0 – 10) onto a continuous variable that behaves like an invader’s growth rate using the function:

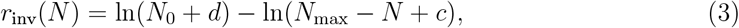

where *N* is the Nugent score, *c >* 0 is a small calibration constant to avoid *ln*(0), and *d* is chosen so that *r*_inv_(*N*_0_) = 0. The parameters *d* = 4.5 and *N*_0_ = 4 were selected by maximizing accuracy and F1-score under Monte Carlo cross-validation. An example of the resulting nonlinear mapping is shown in SI Fig. S3. This log transformation can be interpreted as the inverse of an exponential growth rate, similar to how invasion fitness is analyzed in replicator dynamics.

### Model fitting to data

Returning to our replicator equation growth rate *r*_*inv*_ (Eq.2), we can write it even more compactly as a sum of linear and nonlinear terms:

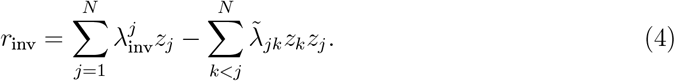

For clarity, we define 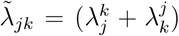 as the coefficient multiplying each product *z*_*k*_*z*_*j*_, to emphasize that interaction effects of species *j* and *k* are captured by one parameter. Looking closely at this expression, it becomes evident that it corresponds to a generalized linear regression with interaction terms, where the dependent variable is *r*_*inv*_ and the independent variables are the resident species frequencies *z*_*i*_,*i* = 1,|..*N*, with Σ_*i*_ *z*_*i*_= 1. With data available for both *r*_*inv*_ and microbiota compositions over many subjects, we can estimate the linear and interaction coefficients using standard regression techniques based on least-squares optimization.

A schematic overview of the modeling approach, from microbiota composition to predicted Nugent score level is shown in Fig. 1.

**Figure 1.**
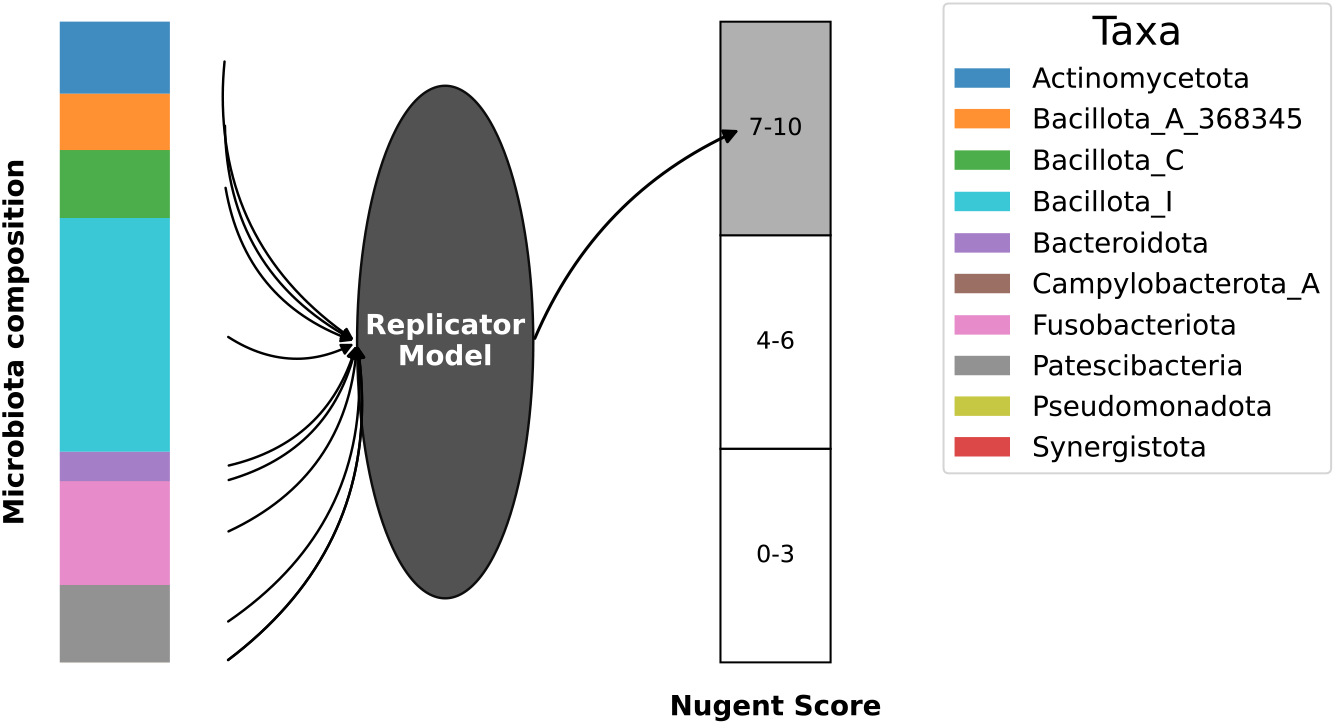
Illustration of model rationale for linking compositional microbiota data to invader growth propensity (Nugent score). Using the replicator equation for estimating microbiota effects on vaginal health status, and onward prediction of vaginal health scores from microbiota composition in new hosts.

In this setting, ordinary least-squares estimation is ill-conditioned and sensitive to sampling noise, likely to produce unstable and inflated coefficients. Thus, due to these limitations and due to the sparsity and high dimensionality of the taxa frequency and pairwise-frequency matrix, particularly for quadratic interaction terms, we decided to fit the model using Ridge regression. Ridge regression introduces controlled regularization that stabilizes parameter estimation while preserving the full model structure.

Parameter estimates were thus obtained by minimizing the regularized objective function

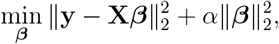

where **X** is the species frequency matrix (containing linear and quadratic variables - species alone *z*_*i*_ and co-occurrence products *z*_*i*_ × *z*_*j*_), **y** is the response vector (*r*_*inv*_-transformed Nugent scores), ***β*** are the model coefficients (*N* linear coefficients and *N* (*N*−1)*/*2 interaction coefficients), and *ε* controls the strength of regularization. When *ε* = 0, we recover the classical Ordinary Least Squares formula.

#### Further modeling choices

We restricted our analysis of Nugent scores to only three clinically relevant categories: healthy (Nugent scores 0-3), intermediate (Nugent scores 4-6) and BV-positive (Nugent scores 7-10). This choice reflects the interpretation and inherent subjectivity of the Nugent scale, and provides additional stability to our algorithm. The middle point of each interval (i.e. 1.5 for healthy samples) was chosen as each class representative.

For the microbiota resolution, we used the *N* = 10 phylum-level groups described in Subsection, and their respective relative abundances, as our independent variables **z**. This results in a total of *N* +*N* (*N*−1)*/*2 = 55 model parameters. Phylum-level classification stabilizes the model fitting by reducing dimensionality while preserving ecological interpretability. For simplicity, we will use the terms phylum and species interchangeably when referring to the *species* within the replicator framework.

#### Preliminary validation tests and parametric optimization

To assess whether this model could reliably recover meaningful ecological parameters, we evaluated its performance on a synthetic dataset designed to mimic key features of the empirical data in [16], with 10-species communities, 3 Nugent score levels and 394 samples. Parameters estimated from the simulated data were compared to their true values using several complementary metrics. With Ridge regularization (*ε* = 0.05), the model recovered 93.5% of linear and 73.5% of interaction coefficient signs, with RMSE values of 0.17 and 0.67, respectively, and correctly predicted health class in 91.7% of samples. Despite imperfect recovery of parameter magnitudes in some cases, these results indicate that the model robustly captures the qualitative ecological roles of microbial taxa and strong predictive performance, supporting application to empirical data (see SI Text S2 for more details).

## Results

With the chosen model-fitting algorithm and its calibrated parameters, we applied the regression framework to the full dataset of 394 women, achieving an *R*^2^ of 72.8%. We first analyzed the obtained ecological parameters and their biological implications (Fig. 2), and how the predictions relate to the original dataset (Fig. 3). Subsequently, we assessed the model’s predictive performance, comparing our estimates to those obtained via standard machine-learning approaches. For this, we used a Monte-Carlo cross validation approach using one subset of the data for parameter estimation and another for prediction of Nugent scores. In our evaluations, we considered both the case of 3 categories for vaginal health state, as well as a binary classification distinguishing only BV-negative (healthy and intermediate; Nugent scores 0-6) and BV-positive (Nugent scores 7-10) samples.

**Figure 2.**
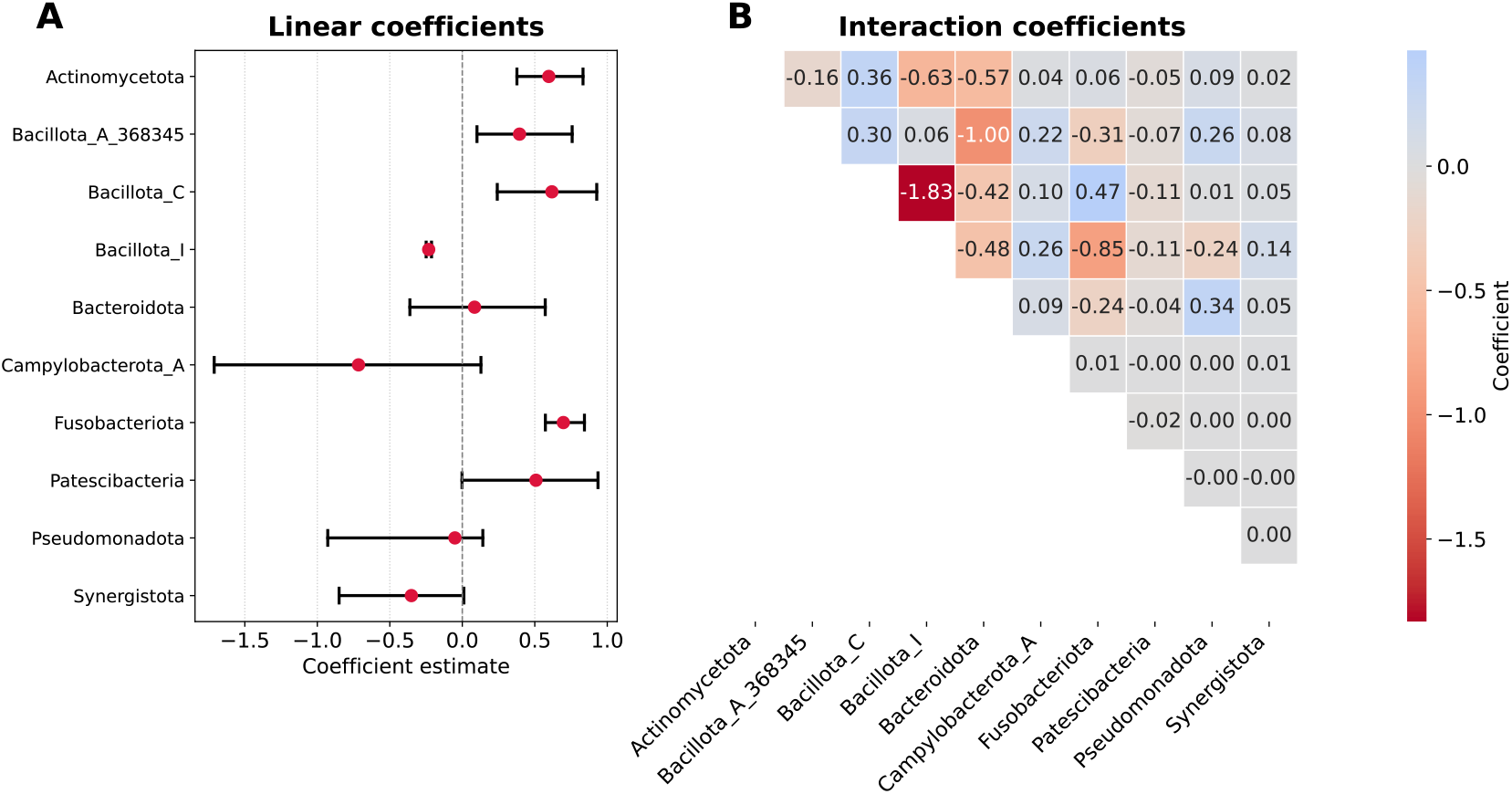
Estimated linear (A) and interaction (B) coefficients for the effects of vaginal microbiota on BV risk, obtained using Ridge regression with *ε* = 0.05. The confidence intervals for the linear terms were generated through a bootstrap method (see SI Methods). The confidence intervals for the interaction terms are available in the SI (SI Figs. S4, S5).

**Figure 3.**
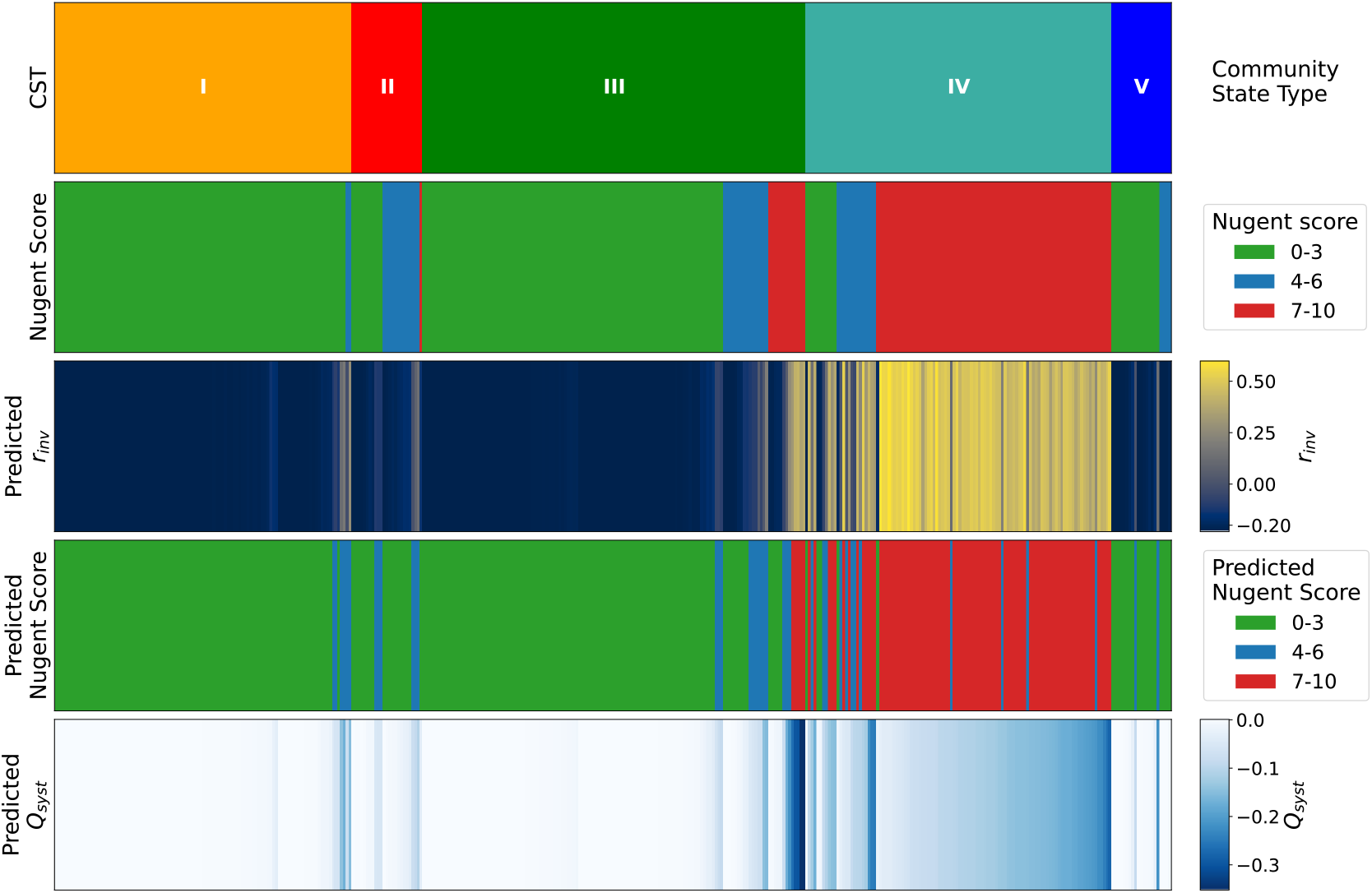
Comparison between empirical structure and model-derived predictions across community state types (CSTs). The first two bars show women’s Nugent scores organized by CST, reflecting the structure of the original dataset [16]. Bars 3–5 display the modeled quantitites in our framework, namely: predicted *r*_*inv*_, retransformed Nugent score predictions, and the replicator system’s invasion resistance metric, *Q*_*syst*_.

### Microbiota species and pairs impact differently vaginal pathobiont invasion

The influence of each species and species pair on the transformed Nugent score is summarized in Fig. 2. This figure illustrates the direct effect of each species and species pair on vaginal health. Linear coefficients (Fig. 2A) indicate how the typical invader invades each species in the system, and the interaction coefficients indicate how the resident species pairs jointly affect the invader’s growth rate (Fig. 2B). Notice that, within this framework, a negative coefficient in A and a positive coefficient in B correspond to a healthier vaginal microbiome.

#### How individual species affect BV risk

The phylum with the strongest negative association with the outcome is *Bacillota_I*, which includes members of the *Lactobacillus* genus, particularly *L. crispatus, L. gasseri, L. jensenii*, and *L. iners*. It is the only coefficient entirely in the negative part of the axis and exhibits the narrowest confidence interval. This finding is consistent with extensive evidence demonstrating the protective role of these species in maintaining a stable and healthy vaginal environment and in reducing the risk of dysbiotic community shifts [30–34].

In contrast, phyla such as *Actinomycetota, Bacillota_A_368345, Bacillota_C*, and *Fusobacteriota* exhibit positive coefficients. These groups include genera and species such as *Sneathia vaginalis, Sneathia sanguinegens, Megasphaera, Gardnerella vaginalis*, as well as members of the class *Clostridia* and the genus *Prevotella*, which have previously been associated with adverse vaginal health outcomes [35–38].

*Synergistota, Campylobacterota_A, Pseudomonadota*, and *Patescibacteria* appear near the center of the coefficient distribution. Although the first three show negative associations and *Patescibacteria* a positive association, the bootstrapped confidence intervals suggest that these effects may not be statistically robust. Furthermore, because *Synergistota* and *Campylobacterota_A* were present at very low abundances in the cohort, their estimated effects should be interpreted with caution.

#### How species interactions affect BV risk

If we now examine Fig. 2B, we observe that most interaction coefficients are close to zero. The main exception involves terms associated with *Bacillota_I* and *Bacteroidota*. With the exception of *Bacillota_A_368345*, all interaction coefficients related to *Bacillota_I* exceed 0.1 in absolute value. The strongest interactions are observed with *Bacillota_C, Fusobacteriota, Actinomycetota*, and *Bacteroidota*. Given the dominance of *Bacillota_I* (Lactobacillus) in the vaginal microbiome, these interaction terms contribute substantially to the effective invasion dynamics. As an example of the biological structure captured by these coefficients, the negative interactions between *Bacteroidota* (Prevotella) and both *Actinomycetota* (Gardnerella; −0.57) and *Bacillota_A_368345* (Peptostreptococcus; −0.16) mirror observed facilitative relationships among these BV-associated taxa [39–41].

While many of the patterns observed here align with existing knowledge of vaginal microbiome dynamics, two aspects of our results are worth noting. First, these conclusions are derived entirely from a mathematical framework that integrates only static snapshots of vaginal communities and their corresponding Nugent scores, demonstrating that meaningful biological information can be extracted from this type of data. Second, we are able to quantify the contribution of each species and species pair to vaginal health, providing a mechanistic and quantitative basis for how microbiome composition leads to different health status. Together, these results establish a rigorous, data-driven and mechanistic framework for interpreting vaginal community structure, opening the door to individualized interventions that enable targeted shifts toward healthy regimes based on a quantitative understanding of microbiome–health relationships.

To connect these theoretical insights back to the empirical data, we turn to Fig. 3. Here, we compare the original dataset structure with the model outputs by showing the women’s Nugent scores organized per CSTs, as identified by Ravel et al [16] (first and second bars), alongside the model’s predictions and the quadratic term — predicted *r*_*inv*_, retransformed Nugent scores, and *Q*_*syst*_ (bars 3–5).

### BV risk can be predicted from microbiota composition

Beyond quantifying the signs and magnitudes of the effects of individual species and species pairs on BV risk, our goal was also to test this approach’s predictive performance. For this, we decided to repeat the same regression procedure on randomly chosen subsets of the data, split 80%-20% for estimation/prediction respectively. As a suitable benchmark for comparison, we used several well-known machine learning algorithms. Table 1 compares the proposed mechanistic framework (Replicator) with several classic machine-learning algorithms for classifying microbiota compositions as healthy, intermediate, or BV, as well as in a binary BV-negative versus BV-positive setting. The results show that the Replicator model performs on par with these approaches (82.9% accuracy for three classes and 92.5% for two) while preserving interpretability.

**Table 1:** Comparison of model performance for predicting Nugent score levels from vaginal microbiota profiles. The proposed Replicator model and standard machine learning algorithms (Random Forest [42], XGBoost [43], SVM [44], and KNN [45] - for implementation details see SI Text S3.2) were trained on the same dataset using two- and three-level Nugent classifications. For the Replicator, two-level performance refers to evaluating a model trained on three classes and subsequently assessed using a binary (BV-negative vs. BV-positive) grouping. Reported metrics (mean ± SD) represent averaged accuracy, precision, and F1-scores across Nugent class groups, over 100 independent 80/20 train–test splits (Monte-Carlo cross validation).

| Algorithm | Nr of Nugent levels | Accuracy (%) | Precision (%) | F1-score (%) |
| --- | --- | --- | --- | --- |
| Replicator | 3 | 82.9 ± 3.8 | 81.9 ± 4.6 | 82.0 ± 4.1 |
| Replicator | 2 | 92.6 ± 2.6 | 92.8 ± 2.5 | 92.6 ± 2.6 |
| Random Forest | 3 | 81.2 ± 3.9 | 79.1 ± 4.8 | 79.4 ± 4.2 |
| Random Forest | 2 | 92.5 ± 2.5 | 92.7 ± 2.5 | 92.5 ± 2.5 |
| XGBoost | 2 | 91.5 ± 2.8 | 91.8 ± 2.7 | 91.5 ± 2.7 |
| SVM | 2 | 92.4 ± 2.6 | 92.7 ± 2.5 | 92.5 ± 2.6 |
| KNN | 2 | 92.9 ± 2.7 | 93.2 ± 2.5 | 93.0 ± 2.6 |

## Discussion

In this work, we developed a mechanistic modeling framework that links vaginal microbiota composition to Nugent health scores, providing a new *proof-of-concept* open for future applications. By mapping clinical Nugent scores to the mathematical quantity of *invader* growth rates in a multi-species ecosystem, we propose a slightly more formal approach for vaginal health quantification, placing this medical metric within a well-established replicator-dynamics formulation [22–24].

This enables a detailed quantification of how individual taxa and pairwise interactions between them contribute to vaginal health. Fitting this model to data recovers established ecological patterns, including the beneficial role of *Lactobacillus* species (*Bacillota_I* in our case), while also providing a principled basis for examining the ecological roles of less well-characterized phyla and their interactions. Rather than relying solely on statistical associations, this approach offers a mechanistic interpretation of community structure in terms of fitness, frequency-dependence, interaction structure, and ecological stability.

A key modeling choice was the aggregation of microbial taxa into broader taxonomic groups. This was motivated by the quadratic increase in model parameters with community size, *N* + *N* (*N* − 1)*/*2, which makes inference at finer taxonomic resolution infeasible with the available data. Working at the phylum level represents a compromise between model complexity, biological realism, and interpretability. From a modeling perspective, this aggregation can be viewed as a coarse-graining of the microbial community, where each group represents organisms with broadly similar ecological roles.

Despite this dimensionality reduction, the model retains strong predictive performance, achieving 92.6% accuracy in classifying BV-positive versus BV-negative samples. Structurally, it is important to highlight that the model achieves this even while restricting interactions to only linear and pairwise quadratic terms. While on one hand this constrains the space of possible ecological relationships, excluding higher-order interactions that are recognized as important in microbial ecosystems [46, 47], its high performance shows that lower-order interactions may be sufficient, as they still determine a large part of clinically-relevant phenotypes.

Our method’s performance is comparable to that of several machine-learning approaches addressing the same problem, as reported in [18–21]. In our own experiments, comparable performance is achieved using phylum-level data, both with standard machine-learning models and with the proposed model, despite the use of substantially smaller feature sets (10 phyla rather than up to 50 taxa in some cases). Notably, the proposed model achieves this performance without sacrificing interpretability. Together, these results indicate that a phylum-level representation captures sufficient ecological signal to connect microbiome community structure with Nugent-based assessments of vaginal health, both at a general level and within data stratification.

For example, as reported in [16], there are differences in the compositions of the vaginal microbiome for different ethnicities. Using our model, we were also able to quantify such subtle aspects of the data, and recover similar prediction performance to previous studies employing much higher microbiota resolution [21]. Testing the performance of our model across these different groups (Asian, Black, Hispanic, White), as shown in Table 2, we obtained lower BV risk prediction accuracy in Hispanic women. Although parameters were estimated as shared by all the dataset, the observed differences suggest that allowing microbiota species effects to vary across groups may capture additional structure. Such variation could be implemented in model fitting in different ways, e.g. as typically done with generalized mixed effects models, or be included via nested parameter inference (see [48]). Exploring such extensions would require larger datasets, with potential combination of multiple cohorts.

**Table 2:** Predicting BV risk by ethnicity based on the local replicator. Two-classes were used for subject classification into healthy and unhealthy states. In this analysis, all individuals were fitted jointly, which implicitly assumes that microbial interactions and role are similar across race, age, and other covariates. This choice was made to maximize statistical power and to identify interaction patterns that generalize across the population. The model was trained on a 80/20 split of the data and then its predictive performace evaluated by ethnicity, with two Nugent level groupings (BV-negative and BV-positive).

|  | Asian | Black | Hispanic | White | Overall |
| --- | --- | --- | --- | --- | --- |
| **Accuracy (%)** | 91.8 ± 5.4 | 93.4 ± 5.0 | 89.1 ± 6.4 | 95.4 ± 4.3 | 92.6 ± 2.6 |
| **Precision (%)** | 93.9 ± 5.2 | 93.8 ± 4.9 | 90.4 ± 5.9 | 97.4 ± 3.1 | 92.8 ± 2.5 |
| **F1-score (%)** | 92.4 ± 5.3 | 93.3 ± 5.0 | 89.2 ± 6.3 | 96.1 ± 3.7 | 92.6 ± 2.6 |

Model fitting was performed using ridge regression, motivated by the sparsity of the taxa frequency and pairwise-frequency design matrix, particularly for quadratic interaction terms involving different phyla. In this setting, ordinary least squares can be unstable, with small perturbations in the data leading to large changes in estimated coefficients. Ridge regression stabilizes the fitting process by shrinking coefficients, improving numerical stability and reducing variance [49–52]. This regularization strategy supports robust parameter estimation in high-dimensional ecological models where interpretability of coefficients is essential.

The framework also has certain limitations that arise directly from its modeling assumptions and data structure. Aggregating taxa into broad taxonomic groups, hides strain- and species-level heterogeneity that may be ecologically and clinically relevant. Likewise, inference from cross-sectional community snapshots limits the model’s ability to predict temporal dynamics, feedback processes, and transient states that characterize microbial community transitions. The assumption of shared interaction parameters across individuals and demographic groups further imposes a form of population-level homogeneity that may mask biologically meaningful variation driven by host physiology, behavior, or environment [53]. In addition, the model does not explicitly represent resource dynamics, despite evidence that nutrient availability and metabolic structure may play a causal role in shifting and maintaining vaginal health states [54]. Finally, discretizing Nugent-derived invader growth rates into categorical levels, while necessary for tractable inference, compresses clinical information and may reduce sensitivity to more subtle variations of vaginal health.

These limitations also suggest clear directions for future work. Replicator dynamics provide a natural framework for modeling longitudinal microbiome data [55], and studying how communities evolve and transition between health states. Here, we recover only partial information about the Λ matrix, capturing the net sum of pairwise interactions 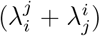 rather than their directional components. As a result, the model does not allow us to understand long-term temporal dynamics, which would require knowledge of each entry of the pairwise invasion fitness matrix between microbiota species. Nevertheless, the results obtained here can serve as constraints for future dynamical models, guiding the development of longitudinal frameworks for vaginal microbiome dynamics, that incorporate temporal data and other factors [53, 56], and enable more detailed prediction of key features.

Ultimately, this framework demonstrates a way of integrating ecological theory with empirical microbiome data and relevant health metrics in a quantitatively interpretable manner. Beyond predictive performance and its immediate clinical application in the BV setting, it provides a practical tool for mapping collective health states to microbiota community composition and interaction profiles, and a foundation for subsequent search of biological mediators underlying estimated model coefficients.

Such a tool can be applied to other within-host environments of clinical or experimental importance such as the nasopharyngeal niche or gut milieu, to explore similar questions of system resilience vs. susceptibility to pathogenic invasion [57, 58]; be it generic, as assumed here, or specific, when a given invading agent is under focus. The quantitative mapping can be further used for precise surveillance and targeted interventions, for example to identify minimal compositional or interaction shifts associated with transitions toward or away from dysbiosis. Although illustrated here as a local phenomenon, using cross-sectional data, the replicator dynamics naturally and readily extends to longitudinal settings, where it may yield deeper mechanistic understanding of microbiota dynamics, including ecological diversity-stability-complexity regimes and their clinical implications.

## Data availability

The bacterial 16S rRNA gene sequences analyzed in this study are publicly available in the National Center for Biotechnology Information (NCBI) Short Read Archive under accession number **SRA022855**. Nugent scores, which serve as an indicator of vaginal health status, were obtained from the original publication that generated the dataset [16].

## Acknowledgements

We thank Sten Madec, Nicola Cinardi, Tomas Camolas and Carlos Andreu for useful discussions in the first stages of this study. This work was supported by Fundação para a Ciência e Tecnologia, Portugal, through project Models4Invasion (FCT grant 2022.03060.PTDC) and project https://doi.org/10.54499/UID/04621/2025.

## Conflict of Interest

The authors declare no conflicts of interest.

## Supporting Information

**SI Dataset**. Vaginal microbiome data with corresponding Nugent scores, Community State Types, and ethnicity annotations.

**SI Text**. Extended methods, supplementary figures, and supplementary tables.

**SI Code**. Source code for data processing, modeling, and figure generation is available at https://github.com/tomasfreire/Modeling_the_human_vaginal_microbiome_replicator

## Notes

### Competing Interest Statement

The authors have declared no competing interest.

