## Supplementary Material for "Modeling the human vaginal microbiome and its protection against pathogens using the replicator framework for invasion"

#### Supplementary Information (SI)

#### S1 Data Processing and Taxonomic Classification

Raw sequence data were processed using the QIIME2 platform (version `[2024.10]` [1]), following standard microbiome preprocessing protocols. Sequences were demultiplexed and subjected to quality filtering based on Phred scores. Amplicon Sequence Variants (ASVs) were inferred using the `deblur` algorithm with a fixed trim length of 200 base pairs.

Taxonomic classification of the resulting ASVs was performed using a pretrained QIIME2-compatible classifier based on the Greengenes2 database [2], which is structured according to the GTDB taxonomy (release 2024.09). Classification was carried out at the phylum level to improve interpretability and reduce dimensionality. In cases where the taxonomic assignment of *Lactobacillus* ASVs was ambiguous at the species or subgroup level, representative sequences were further analyzed manually using the NCBI nucleotide BLAST tool [3] to refine their classification based on sequence similarity.

#### S2 Model validation on synthetic data

We consider a system with $N = 10$ vaginal bacterial species, characterized by pairwise fitness coefficients $\lambda_i^j \in [-1, 1]$ for all $i < j$, and we generate a synthetic dataset consisting of 395 samples (each representing an individual woman) with bacterial relative abundances $z_i \in [0, 1]$, $i \in N$. For each sample, we additionally define an invader species with pairwise fitness coefficients $\lambda_{\text{inv}}^j \in [-1, 1]$, $j \in N$. The invader growth rate $r_{\text{inv}}$ is computed using expression (**??**), with additive Gaussian noise of mean $\mu = 0$ and standard deviation $\sigma = 0.05$, to simulate a misclassification of 1 Nugent score level. The standard deviation was estimated from the empirical distribution of $r_{inv}$ using the 2.5th and 97.5th percentiles, and dividing this range by 11, to simulate the 11 Nugent score classes.

The $N$ invader coefficients $\lambda^j_{\text{inv}}$ (linear terms) and the $N(N-1)/2$ interaction coefficients $\lambda^j_i$ among resident species (quadratic terms) are independently drawn from a uniform distribution $\mathcal{U}[-1, 1]$. For each sample, the bacterial composition vector $\mathbf{z}$ is drawn from a Dirichlet distribution $Dir(1)$.

The continuous invader growth rates are discretized into three levels, mimicking the three Nugent score categories after transformation. We fit the resulting dataset using Ridge regression, varying the regularization parameter $\alpha \in [0, 0.5]$. Predictive performance is evaluated using classification accuracy, while parameter retrievability is assessed separately for linear ($\lambda^j_{\text{inv}}$) and quadratic ($\tilde{\lambda}_{ij}$) coefficients using four complementary error metrics: sign recovery (percentage of coefficients with correctly inferred sign), relative error, absolute error, and root mean squared error (RMSE). Small coefficients were excluded from the last three errors to prevent instability in near-zero coefficients from disproportionately affecting the evaluation metrics.

To capture variability across different systems and avoid bias due to a specific parameter realization, we repeat the entire procedure 100 times, generating 100 independent interaction matrices $\Lambda$ and invader coefficient vectors $\lambda_{\text{inv}}$. Average values across realizations are reported in Fig. S1, with shaded regions indicating one standard deviation.

As expected, increasing $\alpha$ leads to a gradual decrease in $R^2$ and a corresponding increase in mean squared error (MSE), reflecting a loss in goodness of fit due to regularization. However, introducing a small amount of regularization ($\alpha > 0$) substantially improves the recovery of both linear and quadratic coefficients compared to the unregularized case ($\alpha = 0$ - Ordinary Least Squares). As $\alpha$ increases further, parameter estimation errors stabilize and the sign recovery rate for linear coefficients shows no further improvement, indicating diminishing returns from stronger regularization. In contrast, sign recovery for quadratic coefficients deteriorates at larger $\alpha$, related to excessive shrinkage of interaction terms.

Prediction accuracy follows a similar trend: it improves for small $\alpha > 0$ relative to $\alpha = 0$, but eventually decreases as regularization becomes too strong. Based on this trade-off between predictive performance, coefficient stability, and parameter retrievability, we select $\alpha = 0.05$ for all subsequent analyses.

At $\alpha = 0.05$, the model achieves strong predictive performance and partial but meaningful parameter recovery. Averaged across realizations, relative errors are 34% for linear coefficients and 82% for quadratic coefficients, while absolute errors are 0.14 and 0.56, respectively. Corresponding RMSE values are 0.17 (linear) and 0.67 (quadratic). Error metrics were computed after excluding coefficients with small magnitude to avoid inflation of errors, while sign recovery was evaluated on the full set of coefficients. Importantly, the model correctly recovers the sign of 93.5% of linear coefficients and 73.5% of quadratic coefficients. Despite imperfect recovery of coefficient magnitudes, the model correctly predicts the discretized health class (three levels, analogous to Nugent score categories) in 91.7% of simulated samples.

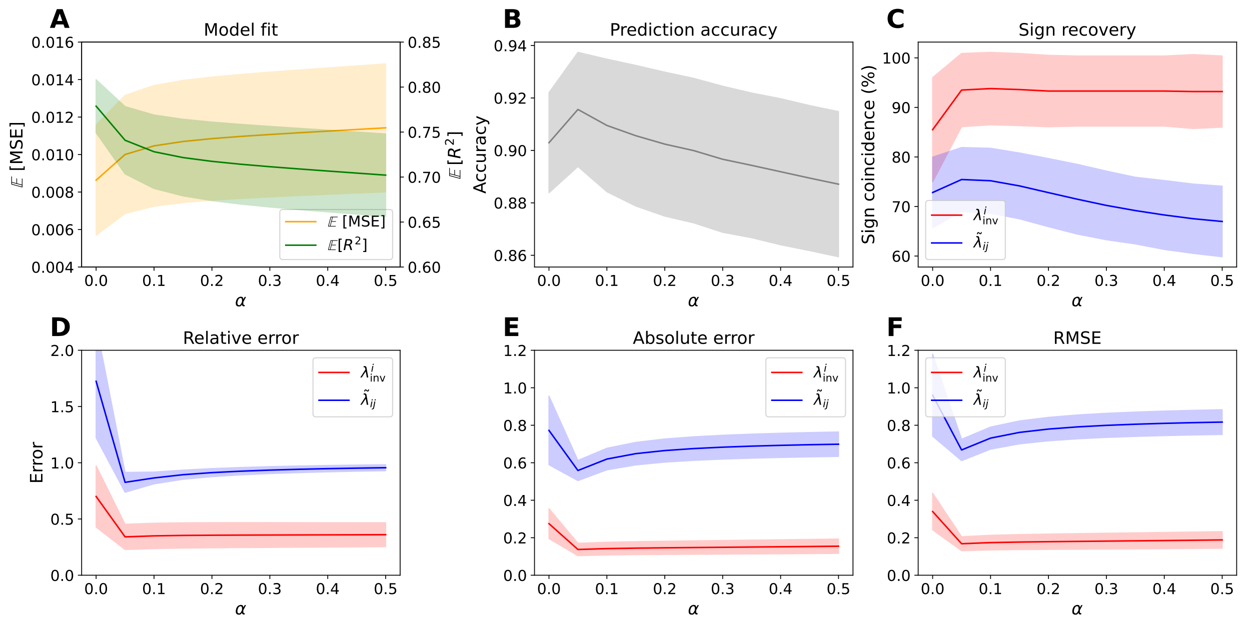


Figure S1: **Results for 100 interaction matrices $\Lambda$ and invader coefficient vectors $\lambda_{\text{inv}}$.** Solid lines indicate mean values across realizations, with shaded areas showing one standard deviation. $\alpha$ is the Ridge regression penalization parameter. **A.** Model fit ($MSE$ and $R^2$); **B.** Prediction accuracy; **C.** Sign recovery of linear ($\lambda^j_{\text{inv}}$) and quadratic ($\tilde{\lambda}_{ij}$) coefficients; **D.** Relative error; **E.** Absolute error; **F.** RMSE.

### S3 Extended Methods

#### S3.1 Estimating confidence intervals via bootstrapping in a Ridge regression

We estimated uncertainty in the model coefficients using a simple bootstrap approach. The dataset was resampled with replacement 1,000 times, and the ridge regression procedure (including the response transformation and feature construction) was repeated for each resample. This produced a distribution of coefficient estimates for every feature. The 95% confidence intervals were then taken as the 2.5th and 97.5th percentiles of these bootstrap distributions. This method is described in [4] for example.

#### S3.2 Details about machine learning implementations

##### Random Forest Implementation

Random Forest classification [5] was implemented using the `RandomForestClassifier` from the `scikit-learn` library in Python. For each iteration, 100 decision trees were trained.

##### XGBoost Implementation

XGBoost classification [6] was implemented using the `XGBClassifier` from the `xgboost` library in Python. For each iteration, 200 gradient-boosted trees were trained with a learning rate of 0.1 and a maximum depth of 6. Feature subsampling was set at 80% for each tree and row subsampling was also 80%.

##### Support Vector Machine (SVM) Implementation

Support Vector Machine (SVM) classification [7] was implemented using the `SVC` class from the `scikit-learn` library in Python. A radial basis function (RBF) kernel was used with a regularization parameter $C = 1.0$ and $\gamma$ set to 'scale'.

##### k-Nearest Neighbors (KNN) Implementation

k-Nearest Neighbors (KNN) classification [8] was implemented using the `KNeighborsClassifier` from the `scikit-learn` library in Python. The model was trained with $k = 5$ neighbors.

#### S4 Performance Evaluation of the algorithms

To evaluate the performance of each method in predicting the Nugent score level, we considered the following metrics:

- **Accuracy:** The proportion of correctly classified samples across all classes:

$$\text{Accuracy} = \frac{\sum_{i=1}^{C}(TP_i + TN_i)}{\sum_{i=1}^{C}(TP_i + TN_i + FP_i + FN_i)}$$

  where $TP_i$, $FP_i$, $FN_i$, and $TN_i$ are the numbers of true positives, false positives, false negatives, and true negatives for class $i$, and $C$ is the total number of classes.

- **Precision (weighted):** The weighted average of precision scores across all classes:

$$\text{Precision} = \sum_{i=1}^{C} w_i \cdot \text{Precision}_i$$

  where

$$\text{Precision}_i = \frac{TP_i}{TP_i + FP_i}, \quad w_i = \frac{n_i}{n}$$

  and $n_i$ is the number of true samples in class $i$, with $n$ the total number of samples.

- **F1 Score (weighted):** The weighted average of F1 scores across all classes:

$$\text{F1 Score} = \sum_{i=1}^{C} w_i \cdot \text{F1}_i$$

  where

$$\text{F1}_i = \frac{2 \cdot \text{Precision}_i \cdot \text{Recall}_i}{\text{Precision}_i + \text{Recall}_i}, \quad \text{Recall}_i = \frac{TP_i}{TP_i + FN_i}$$

These metrics are averaged across the $N = 100$ runs, with a 80-20 train/test split to provide robust estimates of model performance, with standard deviations computed to capture variability.

### S5 Supplementary Figures

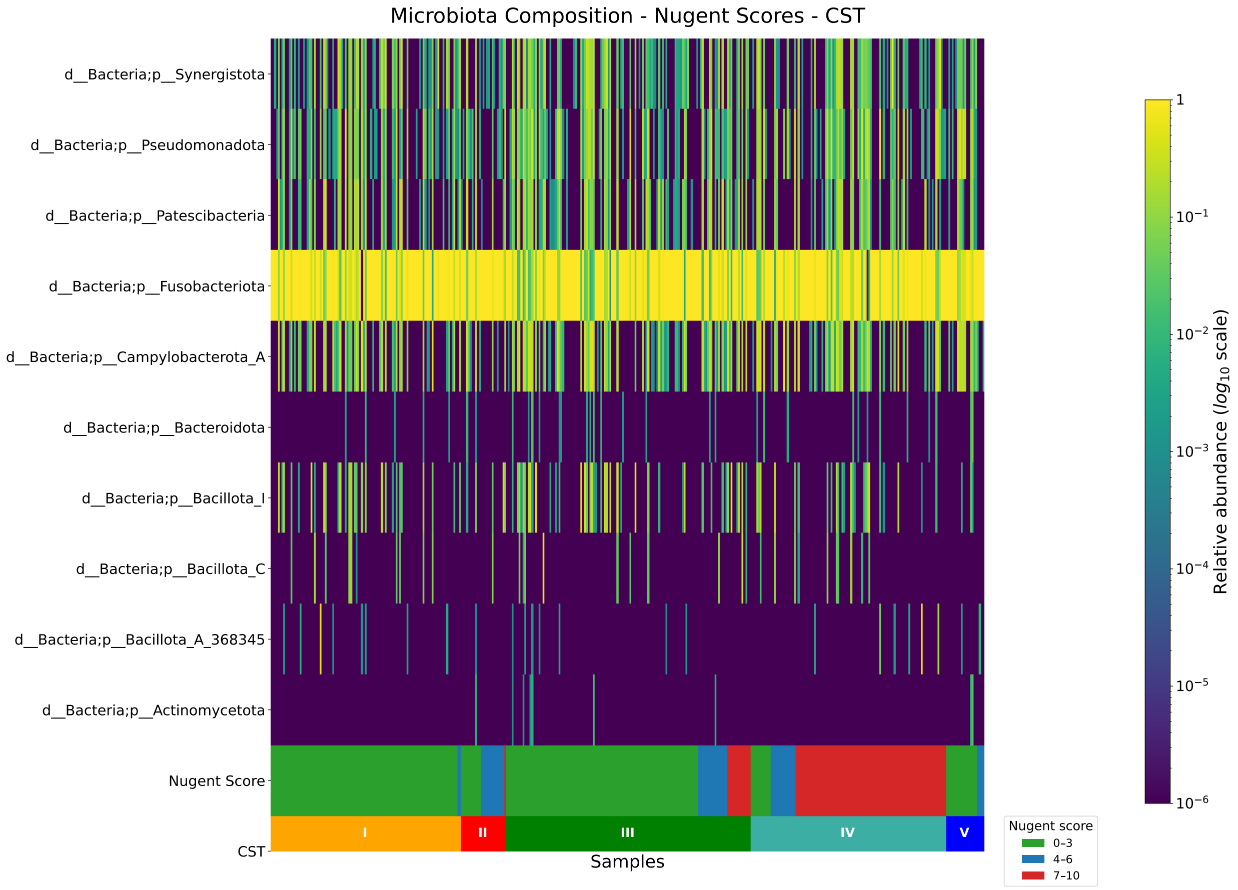


Figure S2: Phylum-level composition of samples. The first ten bars show the relative abundance of identified phyla, while the final two bars denote the corresponding Nugent score and community state type (CST) for each sample.

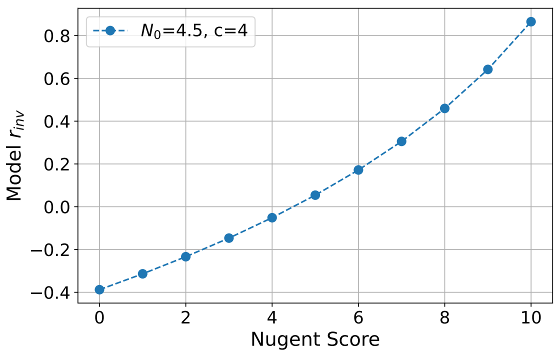


Figure S3: Transformation of Nugent scores to invader growth rates $r_{inv}$, used as the dependent variable in our model fit with microbiota composition data as the independent variables.

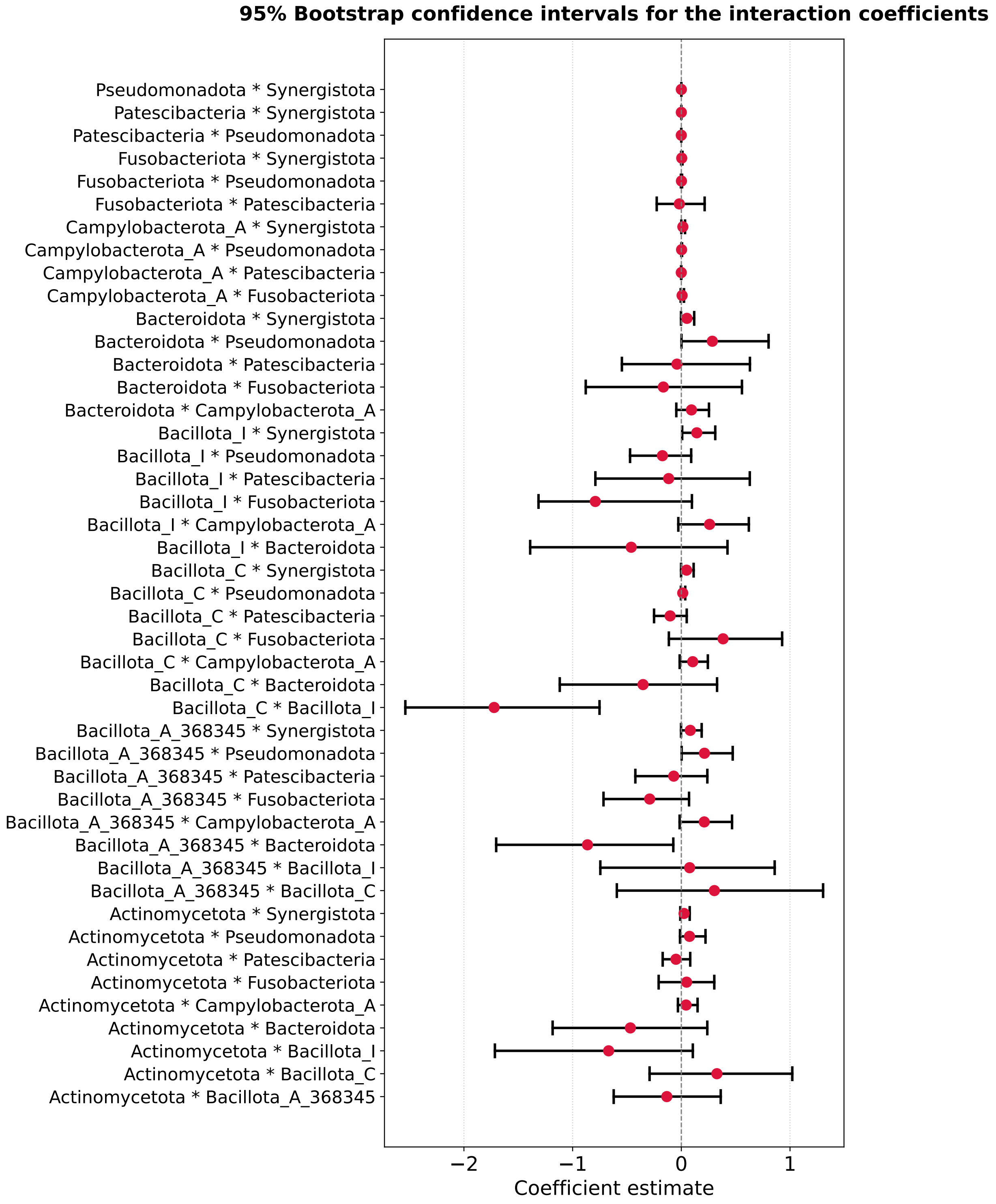


Figure S4: Bootstrapped confidence intervals for the interaction coefficients. Negative terms are associated with an increase in $r_{inv}$.

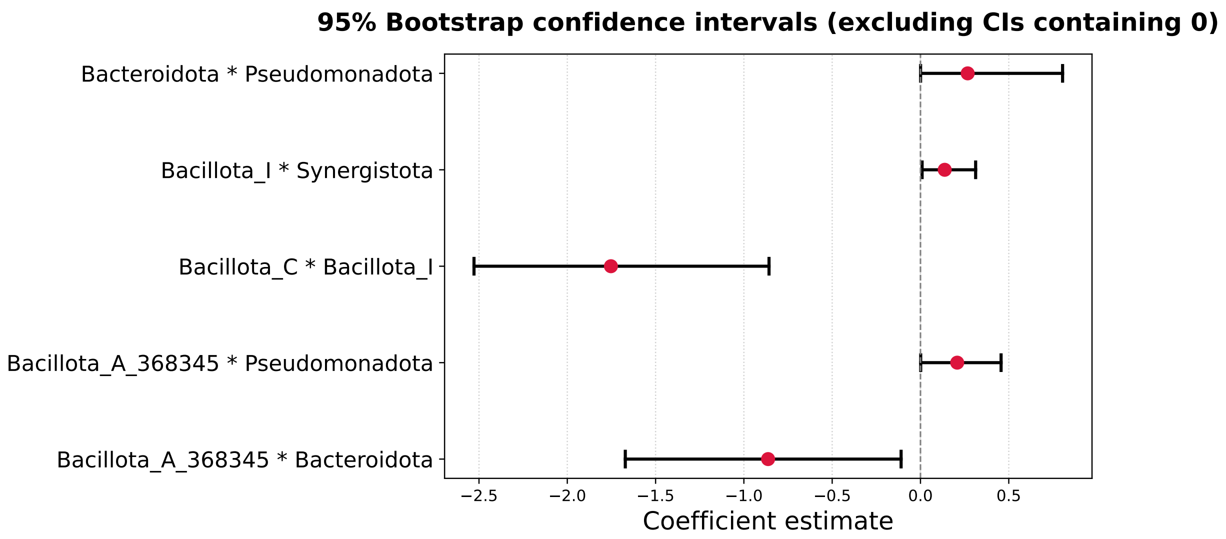


Figure S5: Bootstrapped confidence intervals for the interactions terms, filtered for intervals that don't contain 0.

### S6 Supplementary Tables

Table S1: Representative taxa within each phylum used in this study.

| **Phylum** | **Representative taxa (genera/species)** |
|---|---|
| *Actinomycetota* | *Gardnerella vaginalis*, *Atopobium*, *Eggerthella*, *Mobiluncus*, *Cryptobacterium* |
| *Bacillota_A_368345* | *Roseburia*, *Peptoniphilus*, *Clostridia*, *Finegoldia*, *Anaerotruncus*, *Peptostreptococcus* |
| *Bacillota_C* | *Megasphaera*, *Dialister* |
| *Bacillota_I* | *Lactobacillus crispatus*, *Lactobacillus gasseri*, *Lactobacillus iners*, *Lactobacillus jensenii*, *Aerococcus*, *Streptococcus*, *Gemella* |
| *Bacteroidota* | *Prevotella*, *Porphyromonas* |
| *Campylobacterota_A* | *Campylobacter* |
| *Fusobacteriota* | *Sneathia*, *Fusobacterium* |
| *Patescibacteria* | *Nanoperiomorbus* |
| *Pseudomonadota* | *Sutterella*, *Neisseria* |
| *Synergistota* | *Pyramidobacter*, *Jonquetella* |
